# Some empirical arguments demonstrating that the latent period varies over the course of a plant disease epidemic

**DOI:** 10.1101/148619

**Authors:** Frédéric Suffert, Robin N. Thompson

## Abstract

The latent period is a crucial life history trait, particularly for polycyclic plant diseases, because it determines how many complete monocycles could theoretically occur during an epidemic. Many empirical studies have focused on the variation of the latent period with pathogen or host genotype, or changes in response to environmental factors. The focus on these aspects is unsurprising, as these factors classically form the three parts of the epidemiological triangle. Experiments in controlled conditions are generally used to assess pathogenicity and host susceptibility, and also provide the opportunity to measure the distribution of latent periods in epidemiological systems. Once estimated for one or several pairs of host-pathogen genotypes, the mean value of this important trait is usually considered to be fixed and is often used “as is” in epidemiological models. We show here that the latent period can display non-negligible variability over the course of a disease epidemic, and that this variability has multiple sources, some of which have complex, antagonistic impacts. We develop arguments for four sources of variation that challenge the implicit assumption that the latent period remains constant: daily fluctuations in leaf temperature, nature of inoculum, host stage or age of host tissues, intra-population competition and selection for aggressiveness traits. We focus on the wheat fungal disease Septoria tritici blotch (*Zymoseptoria tritici*), making use of empirical datasets collected during the first author’s own research projects and a targeted literature review. Such empirical epidemiological knowledge is new and potentially important for modelers. While some studies have demonstrated that the distribution of latent periods around the mean value has consequences for epidemiological dynamics, we show that it might also be important for epidemiological modelers to account for changes in this mean value during an annual epidemic. These results may be of critical importance for improving outbreak forecasting.

## Introduction

The latent period is defined as “the length of time between the start of the infection process by a unit of inoculum and the start of production of infectious units” (Madden et al., 2007). It contributes to the generation time of the pathogen, i.e. the length of time in between successive infections, analogously to the age of reproductive maturity of nonparasitic organisms. The importance of the latent period for understanding and predicting pathogen development has long been recognized in plant disease epidemiology (Vanderplank, 1963; Zadoks, 1972). It is a crucial life history trait and component of aggressiveness (Lannou, 2012), especially for polycyclic diseases, because it is one of the major determinants of the number of complete monocycles that could theoretically occur during an epidemic in a single season, which in turn impacts on the final intensity of the epidemic. This view is an oversimplification in the case of pathogens for which disease cycles overlap. Other monocyclic parameters (infection efficiency, infectious period, sporulation intensity) are also important in the adaptive value of a pathogen species and in the predictability of the disease dynamics, but our objective here is not to assess and discuss the relative impacts of these traits on epidemics.

The predictability of disease dynamics depends not only on the ability to assess accurately the mean length of the latent period but also its variability (Cunniffe et al., 2012; Thompson et al., 2016). Ferrandino (2012) clearly showed that the simple use of a population average for the latent period, and also for the infectious period (“the length of time between the start of production of infectious units and the end of production of infectious units”; Madden et al., 2007) is problematic. Using a theoretical model, he demonstrated that the time course of a single annual epidemic does not depend on the average values of the latent and infectious periods alone but is also critically dependent on the variance of these values about their respective means and the covariance between them. This work was followed up by a more detailed analysis showing how the reproduction curves characterizing the production of progeny impact on the speed of an epidemic (Ferrandino, 2013).

Compartmental models used for simulating plant disease epidemics (Kermack & McKendrick, 1927; van der Plank, 1963; Madden et al., 2007) often include an exposed compartment, containing hosts that are in a latent stage (infected but not yet infectious). The length of time that infected hosts spends in this compartment is usually assumed to be exponentially distributed, although other distributions (e.g. gamma distributions) have also been considered in compartmental models (Wearing et al., 2005; Cunniffe et al., 2012; Thompson et al., 2016). These distributions are usually assumed to account for random variation between hosts, rather than systematic differences in latent periods due to e.g. interactions between a pathogen genotype and a host genotype (Viljanen-Rollinson et al. 2005). The mean value of the latent period is also usually assumed to remain constant throughout an epidemic. One exception is the model of citrus greening disease by Parry et al. (2014) in which the mean latent period is assumed to oscillate.

In practice, the length of the latent period changes based on a number of factors. From experimental data, we have identified three origins of variability: (i) experimental uncertainty in the assessment (measurement errors and biases), (ii) phenotypic heterogeneity between individuals within a population (interindividual variance, due for instance to the inherent range of virulence or aggressiveness within the pathogen population), and (iii) variation in the conditions of disease expression, including those due to environmentally induced changes (phenotypic plasticity, whose expression can be amplified for instance by somatic differences in host tissue or differences in microclimate within the plant canopy). Many experimental studies in plant pathology have investigated variations in the latent period with pathogen or host genotype, or its plasticity in response to climatic factors, such as temperature and humidity (Davis & Fitt, 1994; Shaw, 1990; Tomerlin & Jones, 1983; Webb & Nutter, 1997). Such approaches are relevant, because these factors lie at the corners of the epidemiological triangle (host, pathogen, environment; Zadoks, 1972). Most of these studies focused on a mean latent period with statistical features such as standard error (related to the definition of latent period that is used; see below) in order to reduce the uncertainty of the measure, but rarely on the extrinsic variance (i.e. not due to measurement biases, but due to the interindividual variability or the expression of phenotypic plasticity).

There are few empirical data about the time periods over which the latent period of a plant pathogen population changes. However, such a change was detected over pluriannual scales in some cases, for example in poplar rust (Pinon & Frey, 2005). Focusing on soilborne plant pathogens, Leclerc et al. (2014) also noticed that there is little information about how the incubation period (the time between infection and symptom expression) varies temporally in a pathogen population. Interestingly, in that study, the latent period is fixed at zero. A similar observation could be made for the latent period – very few studies consider the possibility that the latent period of a pathogen population may display variability during an annual epidemic.

Our goal here was to highlight key sources of shortterm variability in the latent period, that cause the length of the latent period to vary within a single epidemic season. To this end, we focused on four sources of variability in the development of the wheat pathogen Septoria tritici blotch (*Zymoseptoria tritici*), making use of empirical datasets collected during the first author’s own research projects and a targeted literature review. This fungal disease is particularly suitable for this analysis because the effects of several factors are now well-documented. *Z. tritici* is a cyclic heterothallic pathogen reproducing both sexually and asexually, resulting in infections initiated by two types of spores (ascospores and pycnidiospores), with relative contributions to the epidemic that change over the course of the year (Suffert et al., 2011). The pathogen population displays a high degree of genetic diversity (Linde et al., 2002) and there may be considerable phenotypic variability in the latent period between strains (Morais et al., 2015; 2016). Wheat has a long growth cycle and infections occur from early fall to late spring, under the influence of heterogeneous environmental selective pressures driven by abiotic conditions such as temperature (Lovell et al., 2004a), but also biotic conditions such as the physiological stage of wheat or its fertilization regime (Robert et al., 2005). As Septoria tritici blotch epidemics are polycyclic and result from the integration of many overlapping infection cycles, the latent period is a crucial fitness trait. The latent period is long, facilitating the quantification of any differences by *in planta* experiments and reducing uncertainty in those measurements, and the latent period may display signs of local adaptation to climatic conditions (Suffert et al., 2015). The four drivers of variability in the latent period are described below, and we also demonstrate that accounting for such variability is likely to change pathogen dynamics predicted by mathematical models.

## Operational definitions of latent period can be different

The latent period is regularly measured differently by different plant disease experimenters. This contributes to the variability in the published literature, and makes it impossible to directly compare results. Aligning experimental procedures to allow for direct comparison between experiments might be assumed to be an important first step, and would be advisable in experiments that are similar to each other. However, complete homogenization is neither possible nor desirable. For example, certain definitions are better adapted than others to particular experimental setups because they make it possible to overcome methodological constraints. The latent period for Septoria tritici blotch is usually estimated at the scale of a lesion, as the time between inoculation and the appearance of the first pycnidium (Shearer & Zadoks, 1972; Armour et al., 2004) or, for the sake of convenience, 5% of the final number of pycnidia or 5% of the maximum percentage of area covered by pycnidia (Suffert et al., 2013). Nevertheless, when several lesions rather than a single lesion are considered, particularly if methodological constraints make it necessary (e.g. impossibility of replicating individual inoculation with a given *Z. tritici* genotype using the ascosporic form, contrary to the conidial form; Morais et al., 2015), the latent period is often estimated at the scale of a leaf, as the time between inoculation and the appearance of half of the eventual sporulating lesions (Shaw, 1990; Lovell et al., 2004a). Studies are often conducted by modelers who “search the literature” for experimental parameters to use. We therefore recommend that the operational definition used in a particular experiment is considered by modelers before being used directly.

## The latent period can vary with fluctuations in leaf temperature despite an identical daily mean air temperature

The development of plant pathogens responds strongly to the temperature of the surrounding environment. The effects of temperature are so well recognized in plant epidemiology that linear thermal time (referring to the accumulation of degrees above a given base temperature over a specified period of time; Lovell et al., 2004b) is widely preferred over physical time for assessing and modeling disease development, particularly for Septoria tritici blotch. Consequently, the latent period is usually expressed in degree days rather than as a number of physical days. This accounts, for example, for the decrease in the latent period of *Z. tritici* estimated as a number of days over the spring epidemic period: a 350 degree-day latent period (with a base temperature of -2.4°C; Lovell et al., 2004a) typically corresponds in average to 33 days in early spring (April) but only 22 days in late spring (June) in France (average monthly temperature in Poissy, Yvelines; see https://en.climate-data.org). Taking into account the impact of temperature in this way is however not completely adequate because relationships between temperature and the efficiency or duration of a given epidemiological process are usually nonlinear and often not even monotonic. Consequently, the latent period, while assessed using thermal time, should not be considered constant in time, particularly if the time step used for the calculation is large (e.g. daily), for at least two reasons.

First, thermal time is usually calculated from air temperature, whereas the development of foliar fungal pathogens, including *Z. tritici*, reacts more directly to leaf temperature (the temperature actually perceived by the fungus), which can be very different from air temperature (Bernard et al., 2013). Leaf temperature is harder to measure than air temperature, but it can be estimated indirectly from soil-vegetation-atmosphere transfer (SVAT) models including data recorded at standard weather stations (Xiao et al., 2006).

Second, the latent period is usually assessed under fluctuating temperature regimes, with a thermal 171
scale based on the accumulation of daily mean temperatures. The effects of diurnal fluctuations are, therefore, not taken into account. Bernard (2012) established the impact of two patterns of leaf temperature variation, in which the mean temperatures were equal (18°C) but daily temperature ranges differed (±2°C and ±5°C), on the latent period of *Z. tritici*: the larger temperature range increased the latent period by 1.3 days on average. Similar results have been obtained for other plant pathogens (Scherm et al., 1994; Xu, 1999). The differences in pathogen development between constant and fluctuating environments are partly due to “rate summation” or the Kaufmann effect (Ruel & Ayres, 1999), a mathematical consequence of the nonlinear shape of thermal performance curves (TPCs). The length of the latent period under fluctuating temperatures can be predicted by integrating constant-temperature developmental rates over the fluctuating temperature regime (Hau et al., 1985; Xu et al., 1996):

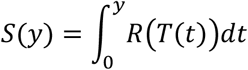

where *S* is the accumulated development over the time interval [0, y], *T(t)* is temperature as function of time *t* and *R*(*T*(*t*)) is the development rate as a function of temperature. *S* is dimensionless and defined as zero initially and one at the completion of a process (i.e. appearance of the first pycnidia).

Finally, degree-hours should be preferred over degree-days (ddpi) once the TPC of the latent period is available.

The mean TPC of the latent period for *Z. tritici* was established empirically, with a limited number of fungal isolates, in natural (Shaw, 1990) and controlled conditions (Bernard et al., 2012). The variability of the latent period among pathogen populations of different geographic origins has never before been characterized in detail. The latent period TPCs presented in Fig. 1a were obtained from two groups of nine *Z. tritici* samples collected from two regions of France with different climates (Brittany and Burgundy). The thermal optimum differed between the two populations: <20°C for the isolates from Brittany, >21°C for the isolates from Burgundy. The effect of temperature on the latent period can differ between pathogen populations expressing local patterns of climatic adaptation.

**Figure 1.**
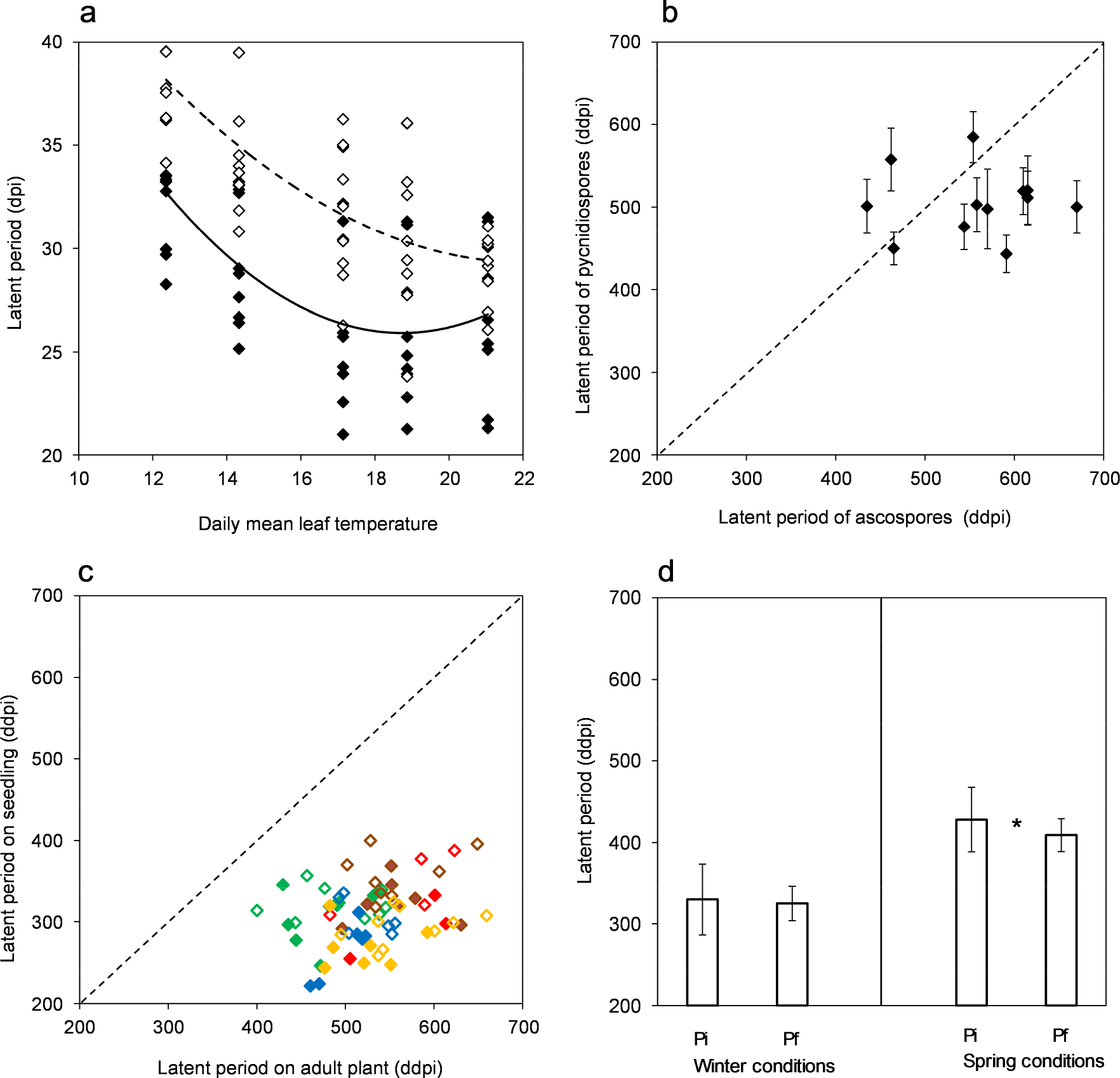
Illustration of four sources of variability in the latent period (expressed in days post-inoculation [dpi] or in degree-days post-inoculation [ddpi] with a base temperature of 0°C) of the wheat pathogen *Zymoseptoria tritici* over the course of an annual epidemic. The latent period was calculated in panels 1a, 1c and 1d as the time between inoculation and the appearance of 5% of the maximum area covered by pycnidia, calculated by fitting a Gompertz growth curve to experimental data as described by Suffert et al. (2013). In panel 1b the latent period was characterized as the time between inoculation and the appearance of 5% of the maximum number of pycnidia in each individual lesion, as described by Morais et al. (2015). **1a.** Effect of the daily mean wheat leaf temperature on the latent period of two *Z. tritici* populations (2 × 9 isolates collected from cv. Apache in two French regions; black diamond = Dijon in Burgundy; white diamond = Ploudaniel in Brittany) assessed after pycnidiospore inoculation on wheat adult plants cv. Apache. The thermal performance curve (the quadratic y = ax^2^ + bx + c, with a =0.17, b = -6.28 and r^2^ = 0.358 for Dijon and a = 0.09, b = -4.10 and r^2^ = 0.548 for Ploudaniel) was adjusted using six replicates per temperature. **1b.** Length of the latent period of 12 *Z. tritici* isolates assessed after ascospore and pycnidiospore inoculation on wheat adult plants cv. Apache (from Morais et al., 2015). Each point corresponds to the mean of several values for pycnidiospore inoculation (vertical bars represent the standard deviation) and a single value for ascospore inoculation. This heterogeneity is due to the impossibility of replicating individual inoculation with a given *Z. tritici* genotype using the ascosporic form, contrary to the conidial form (Morais et al., 2015). Inclusion of replicates for the assessment of the latent period after ascospore inoculation would have made the differences that are currently being displayed far less evident. **1c.** The latent periods of *Z. tritici* populations (2 × 9 isolates collected from cv. Apache in two French regions; colored diamond = Dijon in Burgundy; white diamond = Ploudaniel in Brittany) assessed after pycnidiospore inoculation on wheat seedlings and wheat adult plants cv. Apache for five different wheat cultivars: Apache (green), common to both Brittany and Burgundy; Altamira (red) and Paledor (yellow), mostly cultivated in Brittany; Arezzo (blue) and Altigo (brown), mostly cultivated in Burgundy. Each point represents the mean value from six replicates. The mean latent period was 298 ± 41 ddpi on seedlings and 535 ± 66 ddpi on adult plants. Such differences should be taken into account carefully, especially for multifactorial modeling purposes, as the definition was not the same because of methodological constraints: the ways to assess the latent period (5% of the maximum percentage of area covered by pycnidia) was obtained for adult plants by fitting a logistic model (Suffert et al., 2013) to 17 points but was obtained for seedlings using raw data (5 points) without fitting a model. This difference could explain the low variance in latent period values assessed on seedlings compared to the equivalent measurements on adult plants. **1d.** The mean latent period of two *Z. tritici* subpopulations (2 × 15 isolates collected on seedlings cv. Soissons very early [Pi] in the epidemic and the upper leaf layers at the end of the same epidemic [Pf]), assessed under winter (on wheat seedlings cv. Soissons at 8.9°C) and spring (on adult plants cv. Soissons at 18.1°C) conditions. Asterisks indicate that the mean differs significantly between Pi and Pf (n = 30, *P* = 0.087; from Suffert et al., 2015).

**Figure 2.**
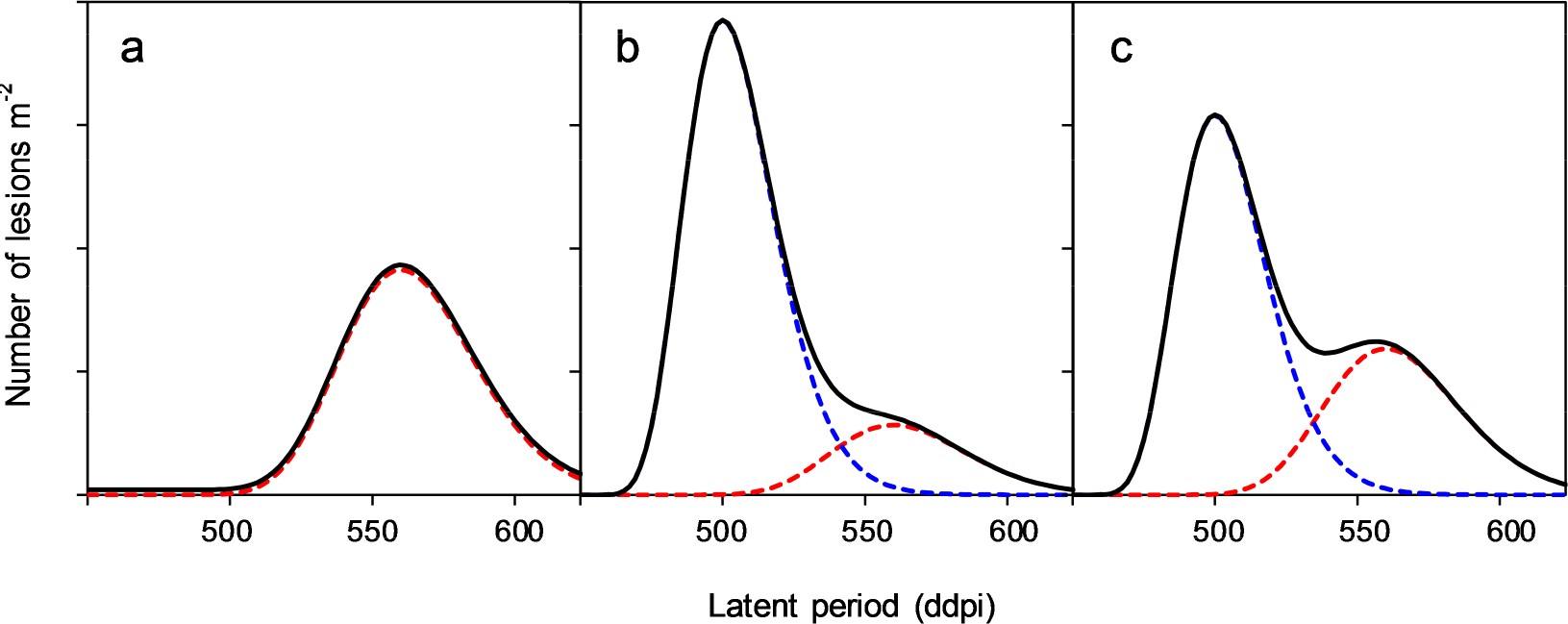
Hypothetical, theoretical distributions of the number of new lesions (on wheat plants, per m^2^ and per week) induced by a pathogen population consisting of different *Z. tritici* strains according to their latent period, taking into account the nature of the spores and the epidemic stage (a = early stage of the epidemic in December; b = intermediate stage of the epidemic in April, c = late stage of the epidemic in June). Red dotted lines correspond to ascospore-initiated lesions; blue dotted lines correspond to pycnidiospore-initiated lesions; solid lines are the cumulative curves. Curves were built with the hypothesis that the mean latent period is 505 ddpi for pycnidiospore infection and 557 ddpi for ascospore infection, based on the results obtained by Morais et al. (2015) and reshown in Fig. 1b. Both latent periods are assumed here to have a gamma distribution, with a similar variance for ascospore and pycnidiospore infections (α = 23 and β = 1.3 for ascospores, α = 11 β = 1.3 for pycnidiospores). The relative heights of the curves are not derived from experimental values, since such experiments do not exist, and should be considered as an approximate order of magnitude. This order of magnitude is inspired by the relative importance of the two types of spores to the epidemic over the growing season found in different studies based on experimental approaches (Hunter et al., 2001; Suffert & Sache, 2011; Duvivier et al., 2013; Morais et al., 2015), theoretical approaches (Eriksen et al.; 2001) or combined approaches (Duvivier, 2015). Overall, it is clear from the literature that the relative role of ascospores vs. pycnidiospores at the end of the season is dependent on the climatic conditions that year. Field reports describing infection of the uppermost (F1) and lowermost green leaves (F3–F4), but healthy middle leaves (F2) (C. Maumené, Arvalis-Institut du Végétal, Boigneville, France, pers. com.), could be explained by ascospore contaminations. Such observations were consistent with both experimental datasets and modeling results. The numbers of ascospores trapped by Eriksen et al. (2001) and Duvivier et al. (2013) suggest that ascospores can play an important role in late infection from mid-April to mid-June. The results of simulations by Eriksen et al. (2001) showed that the proportion of ascospore infection can reach 25% under the most favorable parameter combination. The results of simulations performed by Duvivier (2015), based on three dispersal mechanisms, showed that 50-58% of infections can be explained by wind-dispersed ascospores. The effect of host stage (Fig. 1c), which would likely shorten latent period at the early stage of the epidemic (a), was here not taken into account (see explanations in Fig. 1c).

## The latent period depends on host stage and host tissue age

An increase in the latent period with host development is classically observed for several plant pathogens, such as *Puccinia hordei* (Parlevliet, 1975) and *Puccinia striiformis* (Tomerlin et al., 1983). This finding is consistent with the lack of a univocal relationship between wheat seedling and adult plant resistance. For wheat rusts, for example, many resistance genes are expressed in adult plants but not in seedlings (McIntoch et al., 1995). We assessed the latent periods of two groups of *Z. tritici* isolates collected in two climatically different regions of France (Brittany and Burgundy), on both seedlings and adult plants. We found a large difference between plants of different stages, with a mean latent period of 301 dd for seedlings and 534 dd for adult plants (Fig. 1c). Moreover, other experimental studies have suggested that the susceptibility of wheat tissues varies with leaf layer for synchronous measurements (i.e. on the same date) on adult plants, probably due to differences in leaf age (interactions between the susceptibility of host tissues, natural senescence and nitrogen status; Ben Slimane et al., 2012; Bernard et al., 2013; Suffert et al., 2015). The increase in the latent period length with developmental stage (young vs. adult plants), and, more generally, with leaf age (time between leaf emergence and leaf infection), has been investigated in detail for *Puccinia arachidis* (Savary, 1987). These findings provide further support for our contention that the latent period of a plant pathogen can vary over the course of an epidemic.

## The latent period is strain-dependent and, therefore, affected by competition within a local pathogen population

As mentioned above, the latent period depends on pathogen genotype. Variability within a local pathogen population may be high or low, according to the inherent structure of the population (clonality vs. sexual reproduction that typically leads to high levels of variability). Locally, at the scale of a single annual epidemic, some authors consider average aggressiveness, and, thus, latent period, to be stable (for a given type of spore). Suffert et al. (2015) showed that the mean latent period of *Z. tritici* pycnidiospores can vary significantly during a single annual epidemic: isolates collected on the upper leaf layers of wheat at the end of an epidemic have a shorter latent period than those collected from seedlings very early in the same epidemic. This difference in the latent period between strains, expressed under spring conditions (adult plants, warm temperature) but not under winter conditions (seedlings, cold temperature), suggested that strains with a shorter latent period are selected during the second part of the epidemic (spring), when the disease is propagated by the upward splash dispersal of spores (Fig. 1d). During this period, a short latent period is a key fitness trait conferring a real competitive advantage. These conclusions were corroborated by the significant decrease in between-genotype variance for the latent period. The decrease in the mean latent period of a pathogen population over the course of the epidemic is consistent with the increase in other aggressiveness traits recorded for various fungal pathogens after a few cycles of asexual reproduction (Newton & McGurk, 1991; Villaréal & Lannou, 2000; Andrivon et al., 2007; Le May et al., 2012). Once again, these empirical findings support our key conclusion that the latent period of a plant pathogen can vary over the course of an epidemic.

## TEMPORAL VARIABILITY IN THE LATENT PERIOD IMPACTS ON EPIDEMIC DEVELOPMENT

We have demonstrated that the latent period is likely to vary over the course of a plant disease epidemic, even within a single season. This was the main goal of this article. However, a key question is then whether or not this variability impacts on outbreak dynamics, and consequently whether or not temporal changes in the latent period ought to be including mathematical modeling studies.

We therefore considered the development of a plant disease outbreak at the field scale, both within a season and over the course of multiple seasons (Fig. 3). The model tracks changes in the number of infected sites during the outbreak, where the term “site” refers to a unit of plant tissue that can sustain an infection and further infect other plant tissue (Savary & Willocquet, 2014). In this analysis, we were not intending to replicate the dynamics of successive Septoria tritici blotch epidemics, but rather to test using as simple a model as possible whether or not latent period variation drives pathogen dynamics that are different to dynamics that might be expected if a constant latent period is assumed. Even in our extremely basic model, we found that the number of infected sites each season can be different with variable and constant latent periods. This remained true even when the mean latent period averaged over the season was identical in each case (cf. Figs 3c and 3d). The mean length of the latent period in the (more realistic) variable latent period case might be used in a model with a constant mean latent period if measurements of the latent period are taken at random timepoints throughout a season. As we have shown, however, this will lead to an incorrect representation of disease dynamics compared to using a model in which the length of the latent period varies temporally during the epidemic.

**Figure 3.**
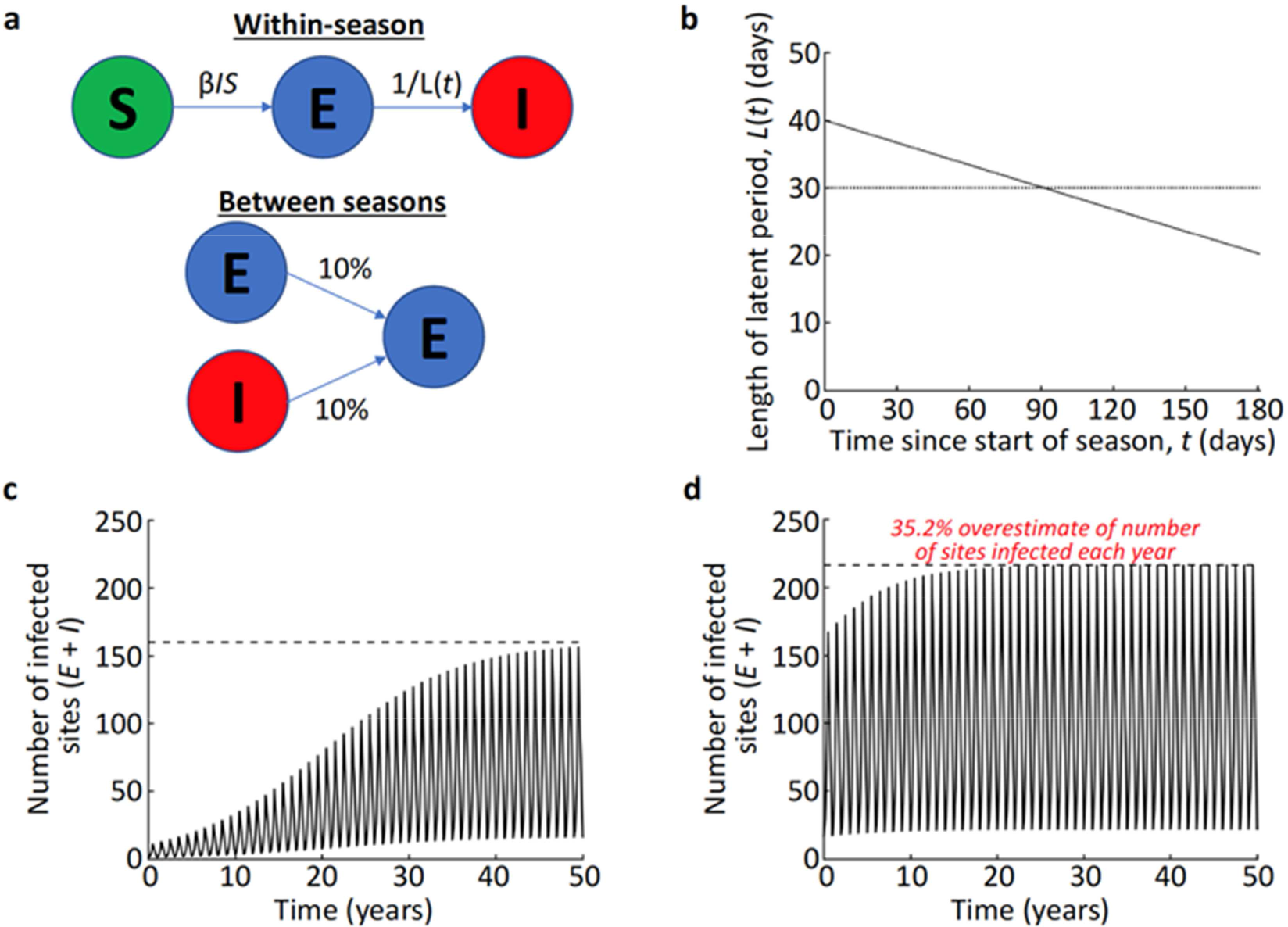
Impact of variability in the latent period on plant pathogen dynamics. We model an epidemic using the classic Susceptible-Exposed-Infected (SEI) model in a host population consisting of *S* + *E*+ *I* = 1000 sites. A site on a leaf can be susceptible (S) i.e. healthy, exposed (E) or infected (I). Between growing seasons, which were considered to last 0.5 years each, there were off-seasons of the same length. In each off-season, 10% of infectious sites from the end of the previous growing season are assumed to found the initial infections (in the *E* class) at the start of the following season. In the case of a disease with dual reproduction modes such as *Z. tritici*, this means that 10% of isolates that induced lesions are then involved in sexual reproduction and generate recombinants that have the capability of causing infections the following season. This proportion is, however, difficult to estimate and so we gave it an arbitrary value, keeping in mind however the case of *Z. tritici* (Suffert et al., 2015; 2018). **3a.** Schematic of the model. Equations for the within-season model are given by d*S*/d*t* = - β*IS*, d*E*/d*t* = β*IS* - (1/*L*(t))*E*, d*I*/d*t* = (1/*L*(t))*E*, where the function *L*(*t*) represents the length of the average latent period of the active pathogen population at time *t* days since the start of the season. In our analysis, we use the infection rate parameter value β = 3 × 10^-5^ per day. **3b.** The latent period lengths that we consider are: i) variable latent period case, *L*(*t*) = 40 - 0.11*t* days (solid black). ii) constant latent period case, *L*(*t*) = 30 days (dotted black); in the variable latent period case, the function *L*(*t*) is chosen so that the mean value is identical to that in the constant latent period case. Specifically, the length of the latent period decreases linearly between 40 days and 20 days, which are values consistent with observed latent periods for *Z. tritici.* **3c.** The number of infected sites, when the latent period varies linearly over the course of the season. The model is simulated over 50 seasons, starting from initial conditions *S* = 999, *E* = 1, *I* = 0. The black dashed line represents the number of infected sites at the end of each season in the long-term when the model settles into regular seasonal dynamics. **3d.** The number of infected sites, when the latent period is constant. The model is simulated over 50 seasons, starting from initial conditions *S* = 999, *E* = 1, *I* = 0. The black dashed line represents the number of infected sites at the end of each season in the long-term when the model settles into regular seasonal dynamics. The number of infected sites at the end of each season is 35.2% greater using a constant latent period than when a potentially more realistic variable latent period is used (*cf.* panel c).

## Discussion

In this article, we have provided empirical evidence to demonstrate that the mean latent period of a plant pathogen population can vary locally, in the shortterm. We have also shown that changes in the latent period can impact on the development of epidemics. Consequently, the mean latent period of the active part of a local pathogen population should not automatically be considered constant over the course of annual plant disease epidemics in future studies.

A significant part of the variability in the length of the latent period is due to the interaction between the interindividual variance within a pathogen population and the expression of its phenotypic plasticity in response to environmental changes; in other words, it is biologically determined. Several empirical arguments justify this assertion as the sources of variation are numerous: daily fluctuations in leaf temperature, nature of the inoculum, host stage or age of host tissues, and selection for aggressiveness traits within a population to name but a few. Some of these sources of variation may have complex, antagonistic impacts. For example, the mean latent period may decrease over the course of an epidemic because of selection for aggressiveness traits driven by biotic or abiotic factors, for instance host stage and temperature (Suffert et al., 2015) in the case of *Z. tritici.* The latent period may also increase at the end of the epidemic due to changes in the ratio of the two spore types resulting from an increase in sexual reproduction before the end of the growing season (Eriksen et al., 2001; Duvivier, 2015). Shaw (1990) suggested that the increase in the latent period that he observed at high mean temperatures reflects the adaptation of *Z. tritici* to local climatic conditions, such as the cool summers in the UK, and a physiological trade-off between an ability to grow rapidly at high temperatures and an ability to grow rapidly at low temperatures. This hypothesis is consistent with the conclusions of Suffert et al. (2015; 2018) that seasonal changes can drive short-term selection for fitness traits, recently confirmed by Boixel et al. (unpubl. data). However, Shaw’s results were obtained in field conditions, and are therefore also impacted on by a number of factors including host stage (the latent period is shorter on seedlings than on adult plants), the use of air temperature rather than leaf temperature (Bernard et al., 2013), and a greater amplitude of daily fluctuations during spring than during winter (Bernard, 2012).

The experimental evidence presented here that challenges the assumption that the mean latent period of a local pathogen population remains constant over the course of an epidemic are original and useful for plant disease experimenters. While the direction (decrease or increase) and the biological causes of these variations are difficult to determine, and accurate characterization of variability in the latent period may require collection of additional data, modelers should consider that the mean latent period may not necessarily take a constant value throughout a plant disease outbreak. Directional variability in the length of the latent period, driven by biophysical processes such as the four sources of variation that we have identified, could be built into epidemiological models.

Sources of short-term variability in the latent period could be analyzed further and potentially incorporated in the case of *Z. tritici* into three types of epidemiological models at least: (i) forecasting models used to simulate the development of annual epidemics and improve wheat protection strategies, e.g. taking into account secondary inoculum pressure to determine the optimal timings for effective fungicide sprays (e.g. Audsley et al., 2005; El-Jarroudi et al., 2009); (ii) mechanistic models used as research tools for understanding the impacts of different epidemiological parameters and processes in driving infectious disease outbreak dynamics, for example discerning the relative importance of pycnidiospore and ascospore infections (e.g. Eriksen et al., 2001; Duvivier, 2015) or the dynamic interaction between plant architecture impacted by cropping practices (nitrogen fertilization, sowing density) and spore dispersal (Baccar et al., 2011); (iii) eco-evolutionary models over several epidemic seasons (Fig. 3) in which the latent period might evolve in response to selective pressures, for example thermal variation (Suffert et al., 2015; Anne-Lise Boixel, pers. com.). There may also be an evolutionary trade-off between intra-and inter-annual scales (Suffert et al., 2018), or an evolutionary optimum driven for instance by the level of nitrogen fertilization (Précigout et al., 2017).

We demonstrated the principle that including directional variability in the latent period, rather than making the common assumption that the mean latent period is constant throughout an epidemic, can change the behavior of a mathematical model (Fig. 3). We chose to consider the simplest possible model in which there is a latent period – namely the SEI model – however of course forecasting would require a more detailed model adjusted for the specific system under consideration. Alterations might include features such as the spatial distribution of hosts and temporal changes in disease management strategies (control of inoculum sources, varietal diversification to limit adaptation of the pathogen population to the host, etc.). For *Z. tritici* specifically, more detailed models exist and could be used (e.g. Elderfield et al., 2017). However, we have demonstrated unequivocally the principle that variability in the latent period should be considered in future modeling studies.

Of course, the values of other epidemiological parameters are also likely to vary temporally (e.g. the infection rate and infectious period). In theory, it might be possible to include variability in those factors, as well as to model complex features in detail such as individual lesion growth dynamics and variability between different leaf layers. However, epidemiological modeling requires some simplifications to be made for tractability, and so that the model can be parameterized. Deciding which factors to include is therefore a challenging balancing act. Here we have shown not only that the latent period can vary, but also that this variation may alter disease dynamics significantly. As a result, considering the impacts of the underlying assumption that the latent period is constant in basic models should be considered further in future work.

Under some circumstances, including detailed descriptions of the latent period may not in fact increase the accuracy of model predictions. Cunniffe et al. (2012) proposed an extension to the generic SEIR model, splitting the latent and infectious compartments and thereby allowing time-varying infection rates and more realistic distributions of latent and infectious periods to be represented. Their results demonstrated that extending a model that has such a simplistic representation of the infection dynamics may not always lead to more accurate results. However, including accurate representations of epidemiological periods in models can be extremely important. Leclerc et al. (2014) conducted experiments on the soilborne pathogenic fungus *Rhizoctonia solani* in sugar beet and used spatially-explicit models to estimate the incubation period distribution. They showed that accurate information about the incubation period distribution can be critical in assessing the current size of an outbreak and the likely efficacy of proposed control interventions.

We have demonstrated that the mean length of the latent period varies over the course of plant disease epidemics, and have identified some sources of variation in the context of *Z. tritici.* Further sources of variability are likely to exist in addition to those considered here. However, we hope that this study will prompt more detailed quantification of the variability in the latent period for a wide range of pathogens, as well as more detailed testing of the circumstances in which this variability should be included in modeling studies. We contend that this will lead to more accurate characterization of pathogen dynamics, which in turn might lead to more accurate disease control. This is of clear plant health importance.

## Acknowledgments

We would like to thank the anonymous reviewers for their valuable suggestions for improving the manuscript. RNT thanks Christ Church, Oxford, for a Junior Research Fellowship.

